# Gappy TotalReCaller for RNASeq Base-Calling and Mapping

**DOI:** 10.1101/000489

**Authors:** Bud Mishra

## Abstract

Understanding complex mammalian biology depends crucially on our ability to define a precise map of all the transcripts encoded in a genome, and to measure their relative abundances. A promising assay depends on RNASeq approaches, which builds on next generation sequencing pipelines capable of interrogating cDNAs extracted from a cell. The underlying pipeline starts with base-calling, collect the sequence reads and interpret the raw-read in terms of transcripts that are grouped with respect to different splice-variant isoforms of a messenger RNA. We address a very basic problem involved in all of these pipeline, namely accurate Bayesian base-calling, which could combine the analog intensity data with suitable underlying priors on base-composition in the transcripts. In the context of sequencing genomic DNA, a powerful approach for base-calling has been developed in the TotalReCaller pipeline. For these purposes, it uses a suitable reference whole-genome sequence in a compressed self-indexed format to derive its priors. However, TotalReCaller faces many new challenges in the transcriptomic domain, especially since we still lack a fully annotated library of all possible transcripts, and hence a sufficiently good prior. There are many possible solutions, similar to the ones developed for TotalReCaller, in applications addressing de novo sequencing and assembly, where partial contigs or string-graphs could be used to boot-strap the Bayesian priors on base-composition. A similar approach would be applicable here too, partial assembly of transcripts can be used to characterize the splicing junctions or organize them in incompatibility graphs and then used as priors for TotalReCaller. The key algorithmic techniques for this purpose have been addressed in a forthcoming paper on Stringomics. Here, we address a related but fundamental problem, by assuming that we only have a reference genome, with certain intervals marked as candidate regions for ORF (Open Reading Frames), but not necessarily complete annotations regarding the 5’ or 3’ termini of a gene or its exon-intron structure. The algorithms we describe find the most accurate base-calls of a cDNA with the best possible segmentation, all mapped to the genome appropriately.

## 1 Introduction and Motivations

To obtain key insights into biological problems – especially, those with important biomedical implications – one may need to observe how a population of cells of heterogeneous types behave over time. By identifying and quantifying the full set of transcripts in a small number of cells at different timepoints and under different conditions, and further aided by sophisticated systems-biology inference tools, the scientists have attempted to fill in the gaps in our understanding of complex biological processes — for instance, those involved in disease progression. In a recent article, entitled: “*Broad Applications of Single-cell Nucleic Acid Analysis in Biomedical Research*,” by Michael Wigler [25] the author discusses the hurdles posed by both the heterogeneity and temporality in cancer as detected by single cell genomic assays that could be easily carried over different stages of cancer progression. A complex picture has emerged from these studies: Namely, that a tumor is a highly heterogeneous mixture of many different cell-types ^1^ and that each cell assumes different cell-states in response to the micro-environment, signaling, metabolic needs with different strategies in different cell-types. Thus an important problem faced by the cancer biotechnologists is that of collecting and interpreting massive amount of transcriptomic data just from a single patient assuming that “in the near future assessing both DNA and RNA content simultaneously from hundreds to thousands of single cells will be quantitatively accurate, as complete as needed, and affordable.”

### 1.1 Challenges of RNA-Seq

In attempting to achieve these goals, one still faces enormous computational and statistical challenges:

**Sequence-based transcriptomic data (‘RNA-Seq’) is fundamentally complex.** (a) Genes can be expressed with widely varying copy numbers that change rapidly, (b) the same gene can have multiple splice variants whose structures remain unannotated and are expressed in unknown and varying proportions, and (c) many genes belong to gene-families sharing high-degree of homology. See [11, 2, 22, 8, 7, 6, 4].
**Short read sequencing technologies (e.g., Illumina HiSeq, etc.) have limitations.** Base-calling errors tend to be rather high for next generation sequencing platforms (more than 1% error in the initial 100bp read, with the error rate rising further with the read-length), which further confounds the analysis of already complex transcriptomic data [9].
**Single-cell RNA-Seq presents additional hurdles.** Firstly, the data quality is lowered by the need for enzymatic pre-amplification. This process significantly truncates the 5’ region of the transcript, resulting in an unavoidable loss of sequence information. Secondly, due to the small amount mRNA present in a single cell at any one time, the number of obtainable reads per cell is much smaller than that obtainable from bulk samples (typically *<* 40 million vs. 150 million+), making rare transcripts harder to detect. See [8, 7, 1, 21, 15].

Existing sequence analysis technologies fail to adequately address these problems [5, 12, 26, 24, 23], significantly limiting the effectiveness of single cell RNA-Seq. A superior base-calling approach, such as the one proposed here, could alleviate the situation considerably, for example, by correctly re-calling ‘poor quality’ bases will effectively ‘salvage’ extra reads that would have been discarded due to low quality. This meaningfully increases the number of reads per run in cases where the sample is of limited quantity (single cells) or is degraded (preserved tissue).

### 1.2 TotalRecaller: Base-calling innovation

TotalReCaller (TRC, [17]) is a rapid base-calling and resequencing platform for NGS (next-generation sequencing), originally created to be versatile in handling various genomics applications. Currently, alternative re-sequencing approaches use multiple modules in a serial pipeline (i.e., without feedback) to interpret raw sequencing data from next-generation sequencing platforms, while remaining oblivious to the genomic information until the final alignment step [5, 12, 26, 24, 23, 17, 3]. Such approaches fail to exploit the full information from both raw sequencing data and the reference genome that can yield better quality sequence reads, SNP-calls, variant detection, as well as an alignment at the best possible location in the reference genome. TRC addressed this unmet need for novel reference-guided bioinformatics algorithms for interpreting raw analog signals representing sequences of the bases (A, C, G, T), while simultaneously aligning possible sequence reads to a source reference genome.

The resulting base-calling algorithm, TotalReCaller (TRC), achieves demonstrably improved performance in all genomic domains, wherever it has been tested. A linear error model for the raw intensity data, coupled with Burrows-Wheeler transform (BWT) and FM-index based alignment create a Bayesian score function, which is then globally optimized over all possible genomic locations using an efficient branch-and-bound approach. The algorithm has been implemented in soft- and hardware [field-programmable gate array (FPGA)] to achieve real-time performance. Empirical results on real high-throughput Illumina data were used to evaluate TotalReCaller’s performance relative to its peers Bustard, BayesCall, Ibis and Rolexa based on several criteria, particularly those important in clinical and scientific applications [17]. Namely, it has been evaluated for (*i*) its base-calling speed and throughput, (*ii*) its read accuracy, (*iii*) its specificity and sensitivity in variant calling and (*iv*) its effect on FRC (Feature-Response Curve) analysis, as used in genome assembly (see [18]).

If our genomic and transcriptomic knowledge was complete and correct (i.e., we have *high quality* references genomes along with its *complete* annotations) then the existing TotalReCaller can derive and use a Bayesian prior efficiently to achieve similar order of high accuracy also in RNASeq applications as in its genomic version [17]. However, more than 50% of the RNA sequences are estimated to be unannotated [22, 7, 4], and complicating the matter, not only are many genes expressed in multiple splice-variant isoforms (whose structures are unknown), but also in cancer, pseudo-genes are often transcribed. These structural variations need to be learned and encoded in the prior used by RNASeqTRC (while allowing for self-index to carry out rapid searches). Two important modifications – one in alignment and the other in data-structures – play a key role in achieving this goal and are described in this paper: namely, (a) branch-and-bound for “gappy” alignment (to reference genome) and (b) a compressed “stringomics” data structure that generalizes BWT to a family of strings (e.g., isoforms). The specific innovative attributes of RNASeqTRC that make it ideal for single cell transcriptomic profiling are summarized:

**High Accuracy.** RNASeqTRC’s empirical Bayesian approach can yield high specificity and sensitivity.
**Robustness Against Incomplete Information.** Encoding the priors by “gappy” references and Stringomics data-structures allows RNASeqTRC to deal with the uncertainty of unannotated genes with no significant loss of performance (compressibility and fast queries).
**High Speed.** RNASeqTRC’s simplicity of structure makes it amenable to hardware acceleration.

## 2 Approach

The proposed approach to transcriptomic assays follows the standard protocols, which have been categorized into the following classes: (1) *Align-then-assemble*, (2) *Assemble-then-align* and (3) *Hybrid Approach* [16]. Since TRC performs simultaneous base-calling and alignment, even when it is used in the *de novo* fashion, it possesses a significant amount of information about the alignments, although this information may vary from transcript-to-transcript. These variations may depend on whether the transcript has been annotated or not, and for unannotated transcript, whether it can be inferred from the reference by a ‘gappy’ alignment. In order to describe the full algorithm precisely and clearly, we have organized the rest of the section, in terms of various building blocks.

### 2.1 Base-Calling without a Reference

The simplest base calling process at the core of TRC involves certain standard pre-processing steps and may vary from technology to technology: for Illumina’s HiSeq technology, we developed linear models addressing crosstalk, fading and cycle synchronous lagging [17]. It mainly uses a dynamic transition matrix in order to filter the raw intensity channels. The model is derived from modeling crosstalk and fading and then extended to include lagging. Since the models are described in great details elsewhere [17], we omit them here.

For simplicity, it is assumed that in each cycle, the sequencing proceeds with one new base at a time (e.g., no lagging in a cycle asynchronous manner). In other words, after the first cycle, there are four possible sequences of length one; after two cycles there are 16 possible sequences each of length two; and after k-cycles there are 4*^k^* possible sequences each of length *k* and so on, which can be represented in a quaternary tree of depth *k*. Among these exponentially many possibilities, a small subset (ideally one unique string represented by a path in the tree) is desired to be identified as the ones very likely to be the correct (or closest-to-the-correct) base-sequence of the DNA. For this purpose, TRC solves a combinatorial optimization problem using Branch and Bound, which statistical estimates the correctness of a solution by an associated score.

The Branch and Bound algorithm [14, 13] is an iterative algorithm based on three consecutive steps. Each cycle performs an iterative process consisting of:

**Branching:** Explore the solution space by adding new leafs to the tree.
**Bounding:** Evaluate the solution space by weighing the leafs of the tree with respect to a suitably chosen *score function*.
**Pruning:** Constrain the solution space by pruning all but the best *b ≤* const, *b ≥* 1 solutions: *b* is the beam-width of the underlying beam-search algorithm. When *b* = 1, note that this is just a *greedy algorithm*. Subpaths of the resulting tree can be augmented with the computed score function, as well as a *p*-value either using a known null-model for the score function or by empirical Bayes method, where null model itself is estimated from the data (e.g., ordering over the score functions of the best *b* solutions computed so far).

Note that an MLE (maximum likelihood estimator) score functions can be computed from the precomputed linear models quite easily using calibrating data (or all the solutions computed so far), without modeling exact chemistry or optimally estimating the parameters of the underlying technology. We recommend a data-driven score-function for this purposes, as it makes the resulting TRC algorithm *technology-agnostic*.

Following the pre-processing step, we may assume that we have a model for following conditional probabilities for the observations: namely, *P_k_*(*X_B_|B*) = conditioned to the underlying base being *B* ∈ {*A, T, C, G*}, it is the probability of estimating the normalized intensity on *B*’s channel to assume a value *X_B_* in the *k*^th^ cycle; *P_k_*(*X_B_|¬B*) = conditioned to the underlying base being *¬B* = {*A, T, C, G*}\ *B*, it is the probability of estimating the normalized intensity on *B*’s channel to assume a value *X_B_* in the *k*^th^ cycle. They may be approximated as Gaussian distributions with the parameters *µ_B_*, *σ_B_*, *µ_¬B_* and *σ_¬B_*:

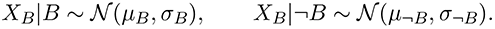

Thus,

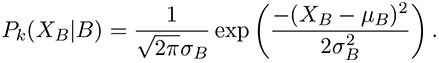

Similarly,

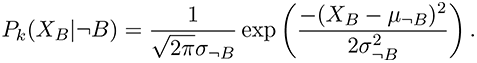

Combining the previous results and computing the log likelihood, we get a score function as shown below:

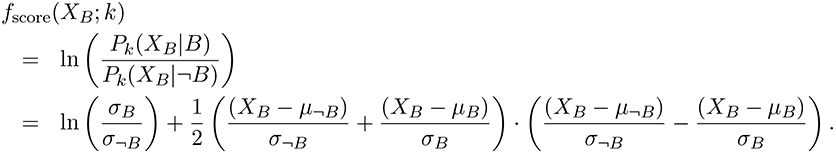

### 2.2 Base-Calling with Gappy Alignment to a Reference Genome

While the approach described earlier, with well-chosen score function extracts as much information as possible to call each base accurately and provides *b*-optimal solutions (*b* = beam-width parameter), ordered according to their scores (or their *p* values or quality scores), it can be further improved in the presence of a Bayesian prior that also provides the marginal probabilities *P_k_*(*B*) and *P_k_*(*¬B*). In the absence of any prior information about the underlying biological system, the most non-informative prior can be chosen to make all *P_k_*(*B*)’s equiprobable for all *B* ∈ {*A, T, C, G*}, taking the value 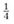 (in which case 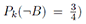^2^; the values can be modified suitably when the *CG*-bias for the reference genome(s) is known, or when the di-neucleotide, tri-neucleotide biases for the reference genome are known (from the reference genome), or when the distribution of *k*-mers over the genome are known. A better solution may be derived from Markov-model of the reference genome (e.g., derived from an estimated HMM), which can be inferred from an assembled reference (genotypic/haplotypic) genome(s), an assembled genome with a single reference along with all the population polymorphisms (e.g., SNP’s, indels, breakpoints, structural variants), or a semi-assembled reference genome with a set of un-phased contigs, or even from just a collection of sequence reads (possibly error-corrected, and organized in a deBruijn graph). A more direct solution can be devised by avoiding pre-processing altogether and simply following a “lazy-evaluation” scheme where *P_k_*(*B*) (and *P_k_*(*¬B*)) are estimated in real-time by aligning the (*k* − 1)-prefix of the sequence, analyzed and ‘called’ so far, to all the locations in the reference genome using efficient compressed and searchable data structures (e.g., BWT, Burrows-Wheeler Transform and FMI, Ferragina-Manzini Index and its variants, see the survey by Navarro and Makinen [20]). Thus the composite score function is:

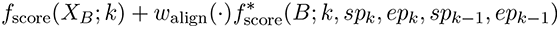

with

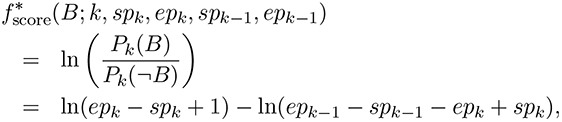

where the FMI’s *sp_k_* and *ep_k_* define the interval in the FMI-dimension corresponding to all the aligned matches in the reference for *B* in the *k*^th^ cycle, which translates in a very straightforward manner to the number of occurrences of the sequences in the reference up to cycle (*k* − 1), which can be calculated by *ep_k_ − sp_k_* + 1. Since the equivalent value after (*k* − 1) cycle is *ep_k−1_ − sp_k-_*_1_ + 1, the corresponding number for “non-matches” to *B* (or matches to *¬B*) is the difference (*ep_k−1_* − *sp_k−1_* − *ep_k_* + *sp_k_*). The estimator can be suitably modified to a “shrinkage estimator,” for instance, one using pseudo-counts, which also avoids various degenerate situations.

It is also straightforward to further generalize the TRC base-callers to more general class of alignments that include “indels,” by simply expanding the 4-character alphabet from {*A, T, C, G*} to a 6-character alphabet {*A, T, C, G, ι, δ*}, where *ι* represents an insertion and *δ* a deletion. Of course, the score function appropriate for a runs of insertion and deletion is more complex, and also requires some amount of “look-ahead” before employing the “pruning” step in the branch-and-bound algorithm. A very naïve way to account for the effect of a ‘gap’ is to introduce another operation *γ*, which indicates that the score function needs to account for a gap in the alignment by restarting a new subtree rooted at a node labeled *γ*. The simplest implementation we describe here lets a new alignment to restart (any where in the genome: the FMI’s being recalculated *ab initio*). In order to avoid trivial gaps, there should be an appropriate gap penalty, and the putative ‘gaps’ will need to be checked (using the FMI’s for substrings between the gaps) in post-processing step. The performance of the ‘gappy’ alignments can be improved significantly, by making sure that the alignment process is sufficiently localized: For instance, in the case of RNASeq applications, it makes sense to limit the alignments only to ORF’s or to run several alignment processes in parallel, with each process using a set of ‘pools’ of ORF’s, where all the ORF’s in the same pool are sufficiently uncorrelated from each other.

However, once such a base-caller is used with priors resulting in ‘gappy’ alignment, the resulting base-calls are expected to be superior to what can be inferred by the traditional base-callers that have been developed for RNASeq applications. But more importantly, from the base call and the ‘gappy’ alignment (the correct one being inferred from the FMI values), one could also infer the locations of exons and splice sites, providing an annotation for the intron-exon structure as well as the splicing isoforms that the data represent.

### 2.3 Base-Calling with Alignment to an Annotated Reference Genome: “Stringomics”

In addition, for RNASeq applications, TRC can also take advantage of the annotated portions of the reference genome, by using a novel data-structure, recently developed by Ferragina and Mishra [10]. In this structure, the exon-intron structures and the multiple splicing-isoforms are encoded efficiently such that the scheme described earlier (for the whole genome) can be extended and generalized easily without sacrificing space and time efficiency. Thus, this “*stringomics*” data-structure supports, as would be expected, the complex topology encoded by the splice junctions connecting groups of exons and is represented as a directed-acyclic graph DAG. Its main function is to align the sequence seen so far as a path in the graph and provides the needed information about the next anticipated base efficiently (e.g., in terms of indices similar to FMI). We sketch the basic ingredients of the “*stringomics*” data structure below, and encourage the reader to consult the full paper [10].

We define a “stringome” to be a family of strings that can be obtained by concatenation of a small number of shorter elemental strings – “stringlets,” which may (or may not) additionally share many common structures, patterns and similarities or homologies. Study of such combinatorial objects have been referred to as “*stringomics*,” as in [10]. The stringomics approach aims to solve various algorithmic problems related to a special case of pattern matching on hypertext. It is built on an underlying graph, which is directed and acyclic (DAG, Directed Acyclic Graph); furthermore, the nodes are assumed to be partitioned into groups, whose strings may have certain additional structures that allow them to be highly compressed.

To be precise, our problem consists of *k* groups of variable-length strings K_1_, K_2_, …, K*_k_*, providing the building blocks for the “stringomes.” The strings are *n* in number, have a total length of *N* characters, and are further linked in a pair-wise fashion by *m* links, defined below more precisely. Each group K*_i_* consists of *n_i_* strings {*s_i_*_1_*, s_i_*_2_*, …, s_in__i_*}, possibly similar to each other. In many situations of practical interest to us, it could be assumed that *|s_ij_| ≤ S*_max_ and *n_i_* is bounded from above by a small constant. The indicator function, 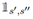 is 1, if there is a link (edge) between the pair of strings (*s’, s”*) and 0, otherwise. It is, then, 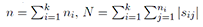 and 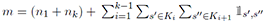. Several complexity bounds can be derived in terms of the parameters *N* and *m*, resorting subsequently to the *k*-th order empirical entropy *H_k_*(K) of the string set K = ∪_*i*_K*_i_* when dealing with compressed data structures [20].

These groups of strings are interconnected to form a multi-partite DAG *G* = (*V, E*) defined as follows. The set *V* consists of *n* + 2 nodes, one node per string *s_ij_* plus two special nodes, designated *s*_0_ and *s_n_*_+1_, which constitute the “source” and the “sink” of the multi-partite DAG and contain empty strings (in order to avoid generating spurious hits). The set *E* consists of *m* edges which link strings of adjacent groups, namely we can have edges of the form (*s*_*ij*′_, *s*_(*i*+1)*j″*_), where 1 *≤ j′ ≤ n_i_* and 1 *≤ j″ ≤ n_i_*_+1_. In addition, the source *s*_0_ is connected to all strings of group K_1_ and the sink *s_n_*_+1_ is linked from all strings in K*_k_*.

The main algorithmic question, to be addressed, is the following: Build an index over *G* in order to efficiently support two basic pattern queries:

**Counting:** Given a pattern *P* [1*, p*], we wish to count the *number occ* of pattern occurrences in *G*.
**Reporting:** Same as the previous query, but here we wish to *report* the positions of these *occ* occurrences.^3^

Various versions of the “Stringomics,” can be created using basic building blocks for: D_K_ (to keep track of the indexing), T_K_ (to organize the underlying strings and stringlets) and P_K_ (to perform 2*d*-range queries in an index-space).

#### Theorem 1

*Listed below are three possible implementations of the “Stringomics” ensemble of data structures, which address three different contexts of use.* ^4^

**I/O-efficiency:** *The following implementation built upon, the String B-tree for* D_K_ *and for* T_K_*, the external-memory Range-Tree for* P_K_*, uses O*(*N/B* +(*m/B*)(log *m/*log log_*B*_ *m*)) *disk pages, which we can safely assume to be O*(*N/B*)*, hence O*(*N* log *N) bits of space*.
**Compressed space:** *The following implementation built upon, the FM-index for* D_K_, *two Patricia tries for* T_K_, *the Range-Tree for* P_K_, *uses N H_k_*(K) + *o*(*N)* + *m* log^2^ *m bits of space*.
**I/O + compression:** *The following implementation built upon, the Geometric BWT for* D_K_*, the String B-tree for* T_K_*, a blocked compression scheme for the strings in* K*, an external-memory Range-Tree for* P_K_*, uses O*(*N* + *m* log *m*) *bits of space*.

We remark that, for various RNASeq applications of immediate interest, any suffix-array like data structure is likely to satisfy our algorithmic needs; nonetheless, we prefer a somewhat more complex implementation based on FM-index as we foresee rapidly growing needs for the technology to scale.

### 2.4 Putting it all together

#### 2.4.1 Base Calling

The RNAseqTRC algorithm works in real-time in the standard manner, but without the fore-knowledge of whether the underlying cDNA (being read currently) corresponds to an annotated gene (in which case the prior is already encoded in the “Stringomics” data structure) or to an unannotated gene, pseudo-gene or a contaminant (in case the prior is available from a possibly ‘gappy’ alignment to the reference genome). Thus TRC runs, in parallel, two (or multiple) branch-and-bound algorithms to call bases with the two sets of priors and compares the resulting score values at the end to decide whether the cDNA examined corresponds to an annotated or unannotated gene.

Additionally, as TRC collects a new dictionary of unannotated genes, it can compile a dictionary of isoforms of genes and pseudo-genes, along with their structural descriptions in terms of exons, introns, and splicing junctions. Periodically, in a “garbage-collection-like” step this dictionary will be examined serially to filter out contaminants (chimeras and sterile transcripts, pseudo-genes, etc.), leaving only the newly discovered genes, rank-ordered by their score functions (or *p*-values). The validated newly discovered genes are then inserted into the existing “Stringomics” data-structure, which will involve modifying the three data-structures: D_K_ (to keep track of the indexing), T_K_ (to organize the underlying strings and stringlets) and P_K_ (to perform 2*d*-range queries in an index-space). The frequency of this “garbage-collection,” step can be determined as the one that optimizes the computational complexity of “dynamization.”

Note that, at this point, the role of TRC can be easily abstracted away (and hence hidden) from the rest of the RNASeq pipeline, as it can treat TRC as just a base-calling module – except that it has the ability to produce better-quality base-calls, and that it can be tuned suitably to take the best advantage of the trade-off between false-positive and negative errors.

#### 2.4.2 Transcriptome Profiling

If our focus was only on the set of transcripts associated with the annotated genes, as would be the case, in many clinical transcriptomic applications, then the simplest strategy would be to keep track of the splice-junctions (i.e., the edges in the Stringomics graph) corresponding to the reads seen from the entire set of reads. The paths in the Stringomics data-structure induced by the edges, labeled by the tracking of splice-junctions, correspond to the splice-variants isoforms, and a rough estimate of such paths can be inferred by a max-flow algorithm running on the graph. However, a much better estimate for the expressed transcripts and their copy number can be obtained from a Bayesian algorithm that, in its prior, models the distributions of the data that correspond to a particular hypothesized transcriptomic profiling.

#### 2.4.3 Transcriptome Assembly

In certain applications, in addition to transcription profiling, it would be necessary to discover mutational changes to transcripts, transcript-editing, new transcripts, new splice-variant isoforms of known/annotated transcripts, or even sterile transcripts (e.g., resulting from pseudo-genes). For such applications, the reads would need to be accurately assembled, which is complicated by the readlengths, quality of base-calling, and various subtle statistical issues, related to variable coverage, estimation of optimal parameters, strand-specificity, etc. The advantage provided by RNASeqTRC are manifold: (1) base-calling accuracy, (2) longer reads, (3) information from alignments to stringomics and reference (that are stored by FM-indices or D_K_/P_K_ structure in Stringomics). These information provide important ingredients to check local correctness of the string-overlaps, and can be summarized by a global score function. Overlap-Layout-Consensus-based global-optimizing algorithm, such as SUTTA [19], can be used with these information to assemble the reads and count the coverage in each transcript-assembly to create a transcriptional profile for all transcripts (sterile or otherwise), and to discover those assemblies that fail to match any of the known annotated transcripts (or fail to align to the reference by a ‘gappy’ alignment).

As discussed earlier, the strategies for whole genome transcript-analysis are usually categorized in terms of three related approaches: (1) Align-then-assemble, (2) Assemble-then-align, and (3) Hybrid [16]. In terms of these categories, the approach described here would be considered a hybrid approach as the underlying base-caller, TRC, automatically aligns to all the known information, such as references, annotations and variations (provided in its prior), and uses these information in basecalling, assembly, validation and discovery.

## 3 Conclusions

This paper initiates the study of transcriptional analysis using very accurate and efficient algorithms, that can be eventually implemented in hardware to run in real-time. Our algorithm efficiently uses Bayesian priors to improve accuracy, and since it obtains these priors from the reference genome and its annotations, it would be appropriate to classify it to be a “reference-guided strategy.” As always, the success of reference-guided assemblers depends on the quality of the reference genome being used, but since TRC can optimize the *w*_align_ parameter in its score function, TRC trades off errors (false positives and negatives) in the best possible manner. TRC will not thus be affected very strongly by the “hundreds to thousands of mis-assemblies and large genomic deletions, which may lead to misassembled or partially assembled transcriptomes,” existing in many extant reference assemblies. Another issue, not directly addressed in this paper, arises from certain trans-spliced genes, in which two pre-mRNAs are spliced together into a single mature mRNA, and requires TRC’s stringomics data structure to be complicated further. In the simplest description provided here, such trans-spliced genes (or those with RNA-editing) will show up as uninterpretable new transcripts. Their status: as new discoveries, as chimeras or as contaminants, will have to be determined in a post-processing step.

However, this paper addresses directly the absence of an efficient and reliable algorithm implementing hybrid assembly strategy for short-read transcripts. Martin and Wang wrote [16], recently, “To date, there are no automated software pipelines that can carry out the hybrid assembly strategy. A systematic study is needed to explore which errors are introduced by hybrid assembly approaches. In the align-then-assemble approach, methods need to be developed to detect the errors in the reference assemblies, in order to prevent them from being propagated into the final assembly. In the assemble-then-align approach, measures must be taken to avoid incorrectly joining segments of different genes (i.e., chimeras).” We believe that the proposed approach promises to fill the gap and address the concerns.

## Footnotes

^1^Certain cell-types such as Cancer Stem Cells, Circulating Tumor Cell, Tumor Initiating Cells, appear to be rare, though they assume disproportionately dominant roles in the fate of the tumor.

^2^When these probabilities are equal and constant, they have no effect on the maximum likelihood estimators, and they provide no advantage over the simplest base-caller described earlier.

^3^It is clear that the identification of a pattern occurrence may involve in our DAG setting three integers: one to identify the source string, (optional) one to identify the destination string, and one to specify the offset of the pattern occurrence in the source string.

^4^We note parenthetically that these are not necessarily the best possible combinations but only offer a good trade-off between simplicity and efficiency.

